# Optimization of neuromuscular blockade protocols in cynomolgus macaques: monitoring, doses and antagonism

**DOI:** 10.1101/2023.12.22.573006

**Authors:** Hélène Letscher, Julien Lemaitre, Emma Burban, Roger Le Grand, Pierre Bruhns, Francis Relouzat, Aurélie Gouel-Chéron

## Abstract

**Background:** Neuromuscular blocking agents (NMBAs) are a crucial component of anaesthesia and intensive care. NMBAs are a family of molecules defined by their ability to compete with acetylcholine for binding to the acetylcholine receptor at the neuromuscular junction. This functional homology relies on the presence of ammonium groups in all NMBAs that, however, display vastly different chemical structures. Among animal models, non-human primates (NHP) are an essential model for a great diversity of human disease models but remain poorly characterized for the effectiveness of the diverse NMBAs.

**Methods:** Seven healthy male cynomolgus macaques were randomly assigned to this study. Experiments using macaques were approved by the local ethical committee (CEtEA #44). All animals were anaesthetized according to institutional guidelines, with ketamine and medetomidine, allowing IV line placement and tracheal intubation. Anaesthesia was maintained with isofluorane. Either rocuronium bromine or atracurium besylate was evaluated, with reversal with sugammadex. Monitoring was performed with two devices, TOF-Watch® and ToFscan®, measuring the T4/T1 and the T4/Tref ratios, respectively. Nonparametric Mann-Whitney statistical analyses were done when indicated.

**Results:** NMBA monitoring required adaptation compared to humans, such as stimulus intensity and electrodes placement, to be efficient and valid in Cynomolgus macaques. When administered, both NMBAs induced deep and persistent neuro-muscular blockade at equivalent doses to clinical doses in humans. Rocuronium-induced profound neuromuscular blockade could be reverted using the cyclodextrin sugammadex’s reversal agent. We report no adverse effects in these models by clinical observation, blood chemistry, or complete blood count.

**Conclusion:** These results support the use of non-human primate models for neuromuscular blockade monitoring and testing novel NMBA or their reversal agents.

## INTRODUCTION

Neuromuscular blocking agents (NMBA) are used in anaesthesia to facilitate intubation and surgical procedures. Based on their chemical properties and structures, they can be divided into two families: depolarizing agents, the only molecule of this family being succinylcholine, used for rapid sequence induction, and non-depolarizing agents: aminosteroids molecules (rocuronium, vecuronium, and pancuronium) and benzylisoquinolines (cisatracurium and atracurium). In clinical practice, two significant side effects are related to NMBAs: anaphylaxis (1/10,000 anaesthesia procedures, with fatal outcome in 4%^1^) and absence of a full recovery of their paralytic results, called residual neuromuscular blockade (rNMB). rNMB can lead to severe consequences (pneumonia, atelectasis, or prolonged hospital stay^2,3^) and has been reported with an incidence as high as 64% in the post-anaesthesia care unit in patients under non-depolarizing agents without monitoring^4,5^.

Neuromuscular blockade cannot be visually evaluated as no clinical test is sensitive enough. The large inter-individual variability regarding NMBA duration of action prevents the translation of results obtained in one patient to another^6^. It is now stated that “monitoring of neuromuscular blockade intraoperatively is recommended”^7^. Since the 1970s, neuromuscular monitors have been developed to deliver a stimulus to a peripheral nerve to objectively measure the strength of the related muscle contraction. The most common pattern is the Train-of-Four (TOF) stimulation, during which four stimuli are provided at a 2 Hz frequency. This assesses the neuromuscular blockade depth with the number of muscle twitch responses to 4 stimuli, T1 to T4. The intensity of the responses and the T4/T1 ratio indicate the depth of the neuromuscular blockade. Intubation can be performed when the T4 responses disappear. A complete neuromuscular blockade reversal is defined as a T4/T1 ratio above 90%. In clinical practice, experts have recommended with a strong agreement (grade 2+) that “the use of TOF stimulation of the ulnar nerve at the adductor pollicis is probably recommended to monitor intraoperative neuromuscular blockade”.^8^

Non-human primates (NHP) are particularly interesting because their anatomy and physiology, especially of the respiratory system, are similar to humans.^9^ Several translational NHP models have been developed to understand the pathophysiology of respiratory infections, evaluate the efficacy of new anaesthetic protocols, or evaluate the relationship between anaesthetic agents and consciousness loss.^10^ In anaesthesia, NHP models were used to evaluate children’s developmental neurotoxicity associated with short-term or prolonged exposure to general anaesthetics^11^. Anaesthesia and mechanically-assisted ventilation might be required in NHP to develop such preclinical models for chest surgery or in any context that may require mechanically assisted respiration. In these procedures, NMBAs are regularly used in the anaesthetic protocol to control properly animal ventilation^12^. To the best of our knowledge, no published study reported NMBA efficacy and dosing regimens in cynomolgus macaques. Only guidelines can be found from Institutions with NHP animal facilities referring to the type and dosing of NMBA, restricted so far to atracurium, vecuronium, and pancuronium^13^. These guidelines lack, however, indications on potential differences between NHP species (cynomolgus, rhesus, or baboon), on NMBA monitoring procedures, and on the assessment of NMBA reversal by anticholinesterases or NMBA-capture reagents (e.g., sugammadex).

In this study, we define an efficient TOF monitoring method in cynomolgus macaques, assess the kinetics and optimal doses for deep NMB of two non-depolarizing NMBAs (rocuronium and atracurium) and of the rocuronium reversal agent sugammadex. Our results are in agreement with human clinical data and validate cynomolgus macaques as an animal model of choice for monitoring the neuromuscular blockade by NMBA and their reversal.

## MATERIALS AND METHODS

### Non-human primates study design

Seven healthy male cynomolgus macaques (*Macaca fascicularis*) were randomly assigned to this study. All animals were chosen as adult, sexually mature (age > four years), and originating from Mauritian AAALAC-certified breeding centers. All animals were housed within IDMIT animal facilities at CEA, Fontenay-aux-Roses in Bio Safety Level (BSL)-2 facilities (Animal facility authorization #D92-032-02, Préfecture des Hauts de Seine, France) and in compliance with the European Directive 2010/63/EU, the French regulations and the Standards for Human Care and Use of Laboratory Animals of the Office for Laboratory Animal Welfare (OLAW, insurance number #A5826-01, US). Animals tested negative for Campylobacter, Yersinia, Shigella, and Salmonella before being used in the study. Only one gender was used to limit heterogeneity. Experiments using macaques were approved by the local ethical committee (CEtEA #44) and the French Research, Innovation, and Education Ministry under registration number APAFIS#36723-2022041910357437 v1. Animals were clinically followed for three days post-infusion. Clinical examination, body weight, and rectal temperature were recorded at each bleeding time, including a complete blood count on each day and a blood biochemistry evaluation on days 0 and 1. Blood sampling did not exceed 7.5% of the total blood volume per week, following ethical recommendations.

### Animals’ anaesthesia protocol

All animals were anaesthetized according to institutional guidelines. For all procedures, the animals were first sedated with ketamine (Imalgene 1000, 5 mg.kg^-1^) and medetomidine (Domitor, 0.5 mg.kg^-1^). An intravenous (IV) line was placed, followed by tracheal intubation allowing mechanical ventilation (Hallowell EMC Matrix 3002Pro veterinary ventilator), and anaesthesia was maintained with isofluorane (Isoflu-vet 1000 mg/g, 0.5-1.5%). Once complete anaesthesia was assessed, macaque received either rocuronium bromine (Esmeron®, MSD 50 mg/5mL) or Atracurium besilate (Atracurium Hospira, 50 mg/5 ml) by IV bolus infusion. When indicated, animals also received sugammadex (Bridion®, MSD, 200 mg/2mL) for rocuronium-induced neuromuscular blockade reversal. Animals were monitored using a Nihon Kohden® PVM-2703 monitor during all the anesthesia procedures. Physiological parameters (capnography, heart rate, respiratory rate, oxygen saturation, arterial blood pressure, and temperature) were followed in real-time and recorded in a chart every 10 minutes. Neuromuscular blockade monitoring devices were installed before NMBA infusion. Mechanical ventilation was maintained for the whole procedure until complete recovery of the neuromuscular function, assessed with a TOF ratio > 90%. All experiments were performed under the supervision of a trained anaesthesiologist and a veterinarian. After the procedures, anaesthesia was reversed with atipamezole (Antisedan, 0.5 mg.kg^-1^), and animals were resumed to their cage under supervision until complete recovery.

### TOF monitoring protocol

Two different monitors were simultaneously used to control and validate the neuromuscular blockade induced in macaques: TOF-Watch® (Alsevia pharma®, Paris, France), which measures T1 to T4 values and calculates the T4/T1 ratio, and ToFscan® (Idmed®, Marseille, France) that additionally calculates the T4/Tref ratio which is a reliable marker of neuromuscular blockade recovery and allows a comparison when the calibration could not be done or when the stimulation is too important^14^. Paediatric hardware was used, when available, to adapt to the macaque’s morphology. A calibration was tried for each device on every animal. Devices were placed on each animal on the ulnar nerve of the left arm for the TOF-Watch® and the right arm for the ToFscan®. The minimal intensity of each device was applied, i.e., 20mA for the ToFscan® and 10mA for the TOF-Watch®. Each software reported each event (injection, loss/resumption of spontaneous ventilation (SV), etc.) simultaneously with a manually filled paper follow-up sheet. Data were extracted through the two dedicated software specific to each monitoring device.

To prevent any movement of the animal during the procedure, the blood pressure cuff was placed on the lower limb of the animal. While the warming pad was maintained, the forced-air warming blanket was stopped during neuromuscular blockade monitoring and restarted as soon as the monitoring ended. The central temperature was monitored throughout the procedure to ensure that no hypothermia occurred, which could have created a bias in the pharmacokinetic parameters of the NMBA used in that experiment.

To ensure the correct placement of the electrodes linked to the TOF devices, the same basic rules that apply to humans were followed^15^. The skin was shaved correctly and cleaned with an alcoholic solution. In human adults, the distance between the two electrodes’ centers should be 3 to 6 cm, with no recommendation in the paediatric setting, which is similar to the size of the macaques used, and we decided to apply a 3cm distance between electrodes. The negative electrode was always placed distally to optimize the response.

As recommended, before initiating the neuromuscular blockade, a stable baseline measure was obtained for both TOF devices for 2 to 5 minutes using the same stimulus pattern (less than 5% variation).

### Statistical analysis

Data were recorded in an internal LIMS and analyzed in Graphpad Prism 9.0 (Graphpad® Software, LLC). Nonparametric Mann-Whitney statistical analyses were done when indicated: *: p<0.05. Results are presented in median with standard deviation or percentages when applicable.

## RESULTS

### Experimental setting

Seven animals were used for the experiments (age 4.9 – 5.6 years, median 5.2 ± 0.21), including blood sampling on days −5, 0, and 1 for complete blood count and blood biochemistry, and on days 2 and 3 only for complete blood count (Figure 1A). Animals were anaesthetized, exposed to NMBA, and monitored on day 0. Some animals were used twice, with a minimum two-week delay between experiments. One animal was excluded as NMBA monitoring could never be assessed (see discussion below). As such, three animals were included in the atracurium group, 3 in the rocuronium group and 4 in the rocuronium+sugammadex group. During neuromuscular blockade monitoring, each device was placed on one limb. Electrodes were positioned above the ulnar nerve (Figure 1B-D) on the anterior lower wrist (Figure 1B-C). Animals were observed for three days following NMBA infusion.

**Figure 1:**
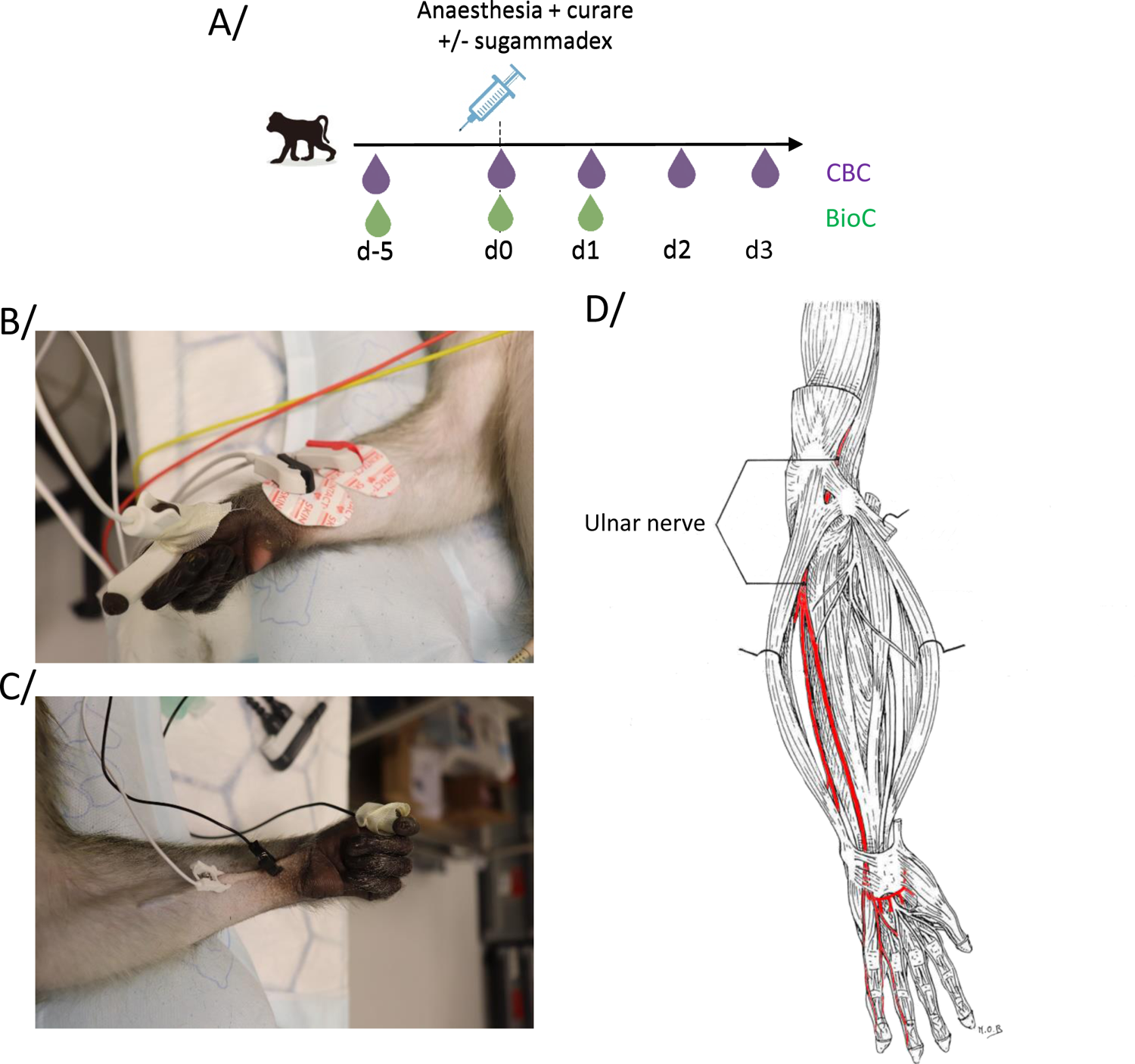
Experimental setting. A/ Animals were clinically followed-up using complete blood count (CBC) and blood biochemistry (BioC) on indicated days. On day 0, blood sampling was done prior to curare infusion. B/ Relative positioning of the positive (red) and negative (black) electrodes to directly stimulate the medial nerve with the Idmed hardware. C/ Relative positioning of the positive (white) and negative (black) electrodes to induce an ulnar nerve direct stimulation with the Alsevia hardware. The accelerometer is positioned on the thumb in each setting (B and C). D/ Anatomical pathway of the ulnar nerve in cynomolgus macaques. Adapted from Marie-Odile Bagnères’ work.

### Accuracy and reproducibility of rocuronium enhanced anaesthesia

Rocuronium-enhanced anaesthesia in cynomolgus macaques was established following a dose-response experiment at 0.2 mg.kg^-1^ of rocuronium intravenous bolus, based on a previous report^16^ (Figure 2). In each case, minimal stimulation was chosen, i.e., 20mA on Idmed® and 10mA on Alsevia® hardware, respectively. Neuromuscular blockade was measured using either T4/Tref (ToFscan®, Idmed®) or T4/T1 (TOF-Watch®, Alsevia®). Loss of SV was observed in all animals about a minute after curare infusion (median 66 seconds ± 7.1). In each animal, loss or deep attenuation of T1 to T4 twitches was observed with either hardware (Figure 2A-F). SV resumption was observed in all animals at a median of 15.9 min ± 3.8 with modest inter-individual variability (Figure 2G). As per recommendation, a 90% T4/Tref (or T4/T1) ratio was obtained for at least 2 minutes before stopping the anaesthesia. A median time of 22.1 minutes (± 11.1) was required to establish two consecutive 90% T4/Tref measures (Figure 2H). Noteworthy inter-individual variability was observed but remained in agreement with the expected variability described in the rocuronium manufacturer specifications (Supplemental). Our calculations were based only on the measures obtained with the TOFscan®, but we found a good consistency between the measurements made with the two devices. The comparison is somewhat challenging as the stimulation intensity between the two devices and the outcome (T4/T1 versus T4/Tref) differ. The TOF-Watch® device (intensity set to 10mA) gave slightly delayed T4/T1 resumption compared to the ToFscan® device (intensity set to 20 mA) with respective medians of 28.2 vs 22.1 minutes, respectively.

**Figure 2:**
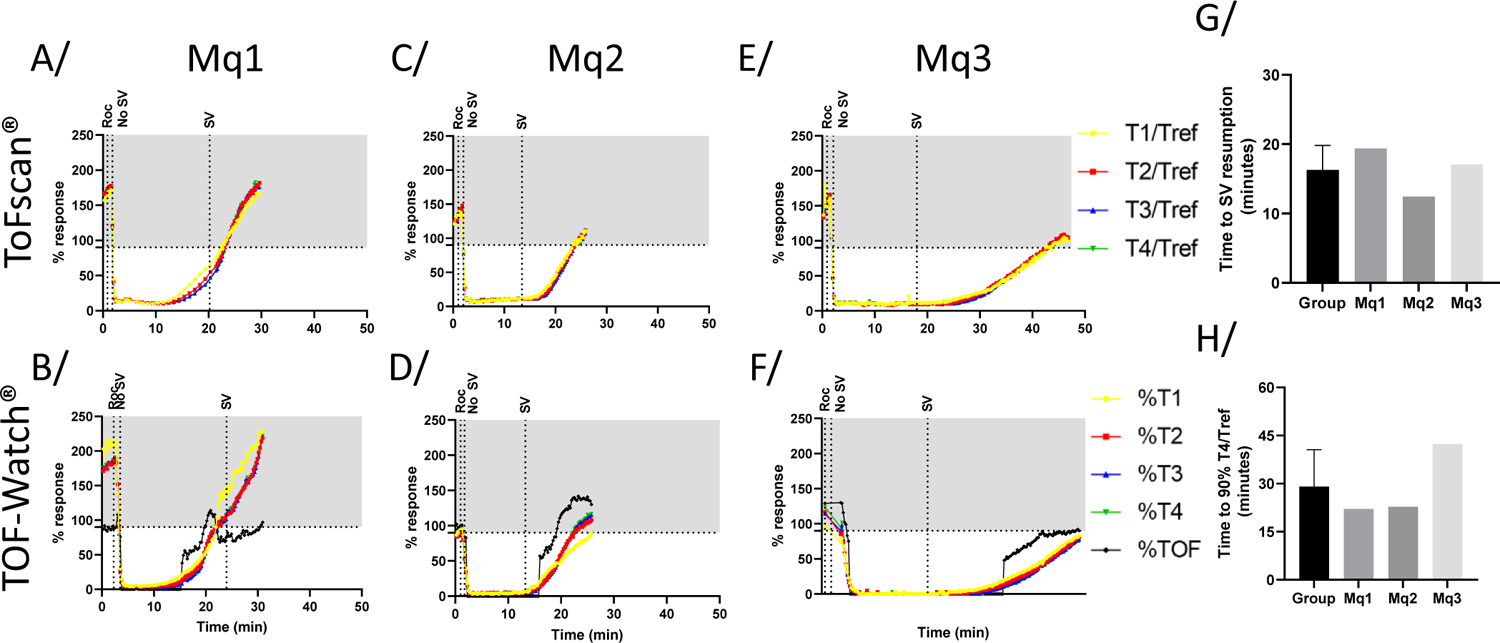
Train-of-four monitoring of three examples of rocuronium-enhanced anaesthesia without rescue. The top line (A, C, and E) are follow-ups done with the ToFscan software (IDMED); the bottom line (B, D, F) were done using the TOF-Watch^®^ software (Alsevia). Regarding the Alsevia software, when no %TOF was given, the values were set to 0. Both were coupled with their respective dedicated hardware. Each column represents a single animal (n=3). G/ Time (in seconds) needed for complete SV resumption from rocuronium injection. H/ Time (in seconds) required to resume at least 90% T4/Tref (either a reference for ToFscan or T1 for TOF-Watch^®^) response classically required to extubate the patient. Roc: rocuronium injection, 200µg/kg dose (i.v). No SV: loss of spontaneous ventilation. SV: Spontaneous ventilation resumption.

### Sugammadex antagonization

After complete neuromuscular blockade was established in four macaques, 1 mg.kg^-1^ intravenous sugammadex was injected to reverse rocuronium-induced neuromuscular blockade (Figure 3). Sugammadex injection induced a rapid and complete recovery in all animals (Figure 3A-D). Compared to spontaneous decurarization, sugammadex permitted a rapid SV resumption and a quicker muscle contractibility restoration (75 seconds ± 22.8 vs 345 seconds ± 277.3, Figure 3E). Neither residual neuromuscular blockade nor adverse events were observed after the sugammadex injection.

**Figure 3:**
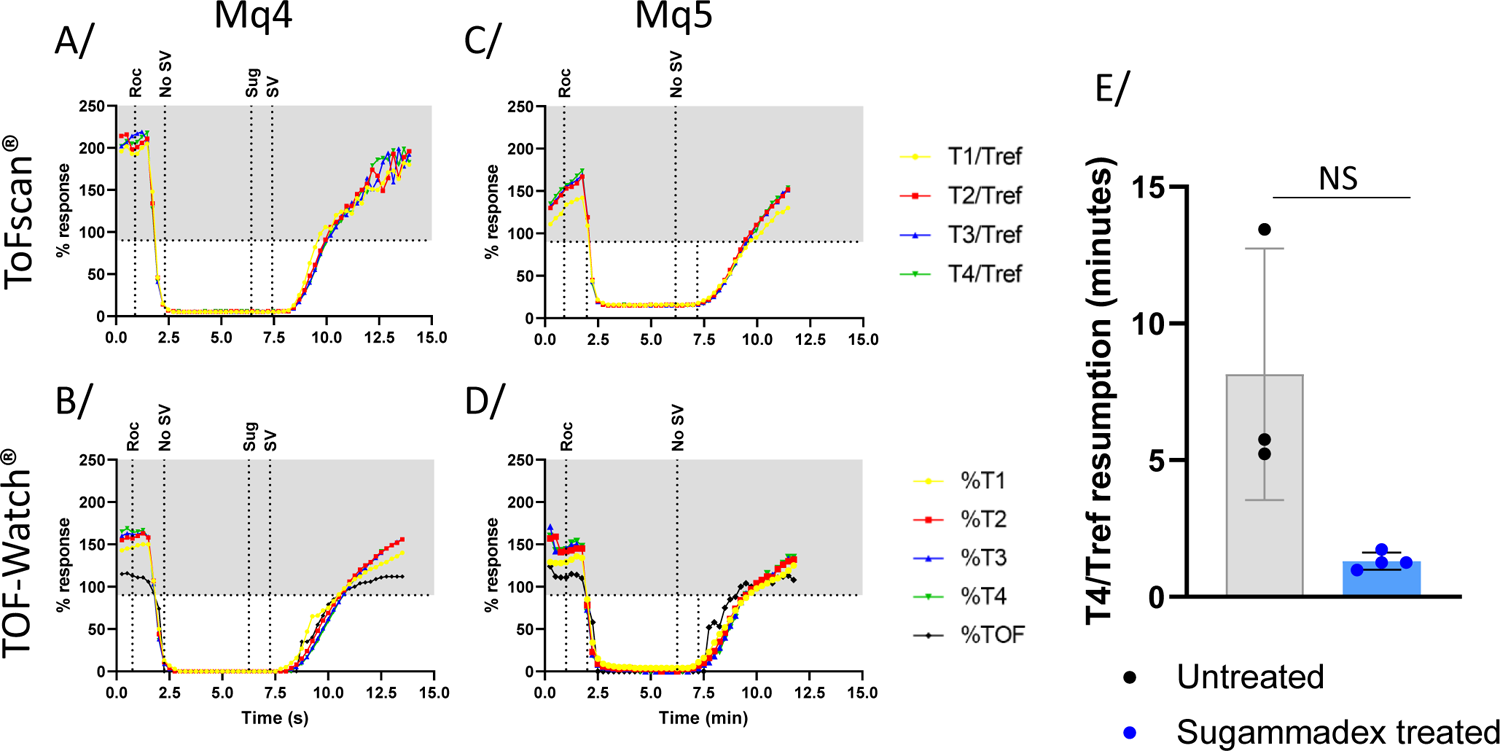
Train-of-four monitoring of two examples of rocuronium-enhanced anaesthesia with Sugammadex rescue. The top line (A, C) is a follow-up done with the ToFscan® software (IDMED); the bottom line (B, D) was done using the TOF-Watch^®^ software (Alsevia). Regarding the Alsevia software, when no %TOF was given, the values were set to 0. Both were coupled with their respective dedicated hardware. Each column represents a single animal (n=4). E/ Time (in seconds) needed for complete spontaneous ventilation resumption from rocuronium injection. For the treated group, this time was calculated between the countermeasure injection and resumption. F/ Time (in seconds) needed to resume at least 90% T4/Tref (either a reference for ToFscan^®^ or T1 for TOF-Watch^®^) response classically required to extubate the patient. For the treated group, this time was calculated between the countermeasure injection and resumption. Roc: rocuronium injection, 200 µg.kg^-1^ dose (i.v). Sug: Sugammadex injection,1 mg.kg^-1^ dose (i.v). No SV: loss of spontaneous ventilation. SV: Spontaneous ventilation resumption.

### Atracurium enhanced anaesthesia

Atracurium-enhanced anaesthesia in cynomolgus macaques was established following a dose-response experiment at 0.4 mg.kg^-1^ of atracurium intravenous bolus^17,18^ (Figure 4). The neuro-muscular blockade was followed-up using the same apparatus and setting as for rocuronium. Loss of SV was observed in all animals after atracurium infusion at a median of 48 seconds ± 8.4 (Figure 4 A-F), non-significantly different from the delay observed after rocuronium injection. SV resumption was observed in all animals at a median of 19.8 minutes (± 3.9), and satisfying neuromuscular blockade reversion (T4/Tref >90%) was noted after 33.8 minutes (± 11.4), with interindividual variability in agreement with the expected variability described in the atracurium manufacturer specifications.

**Figure 4:**
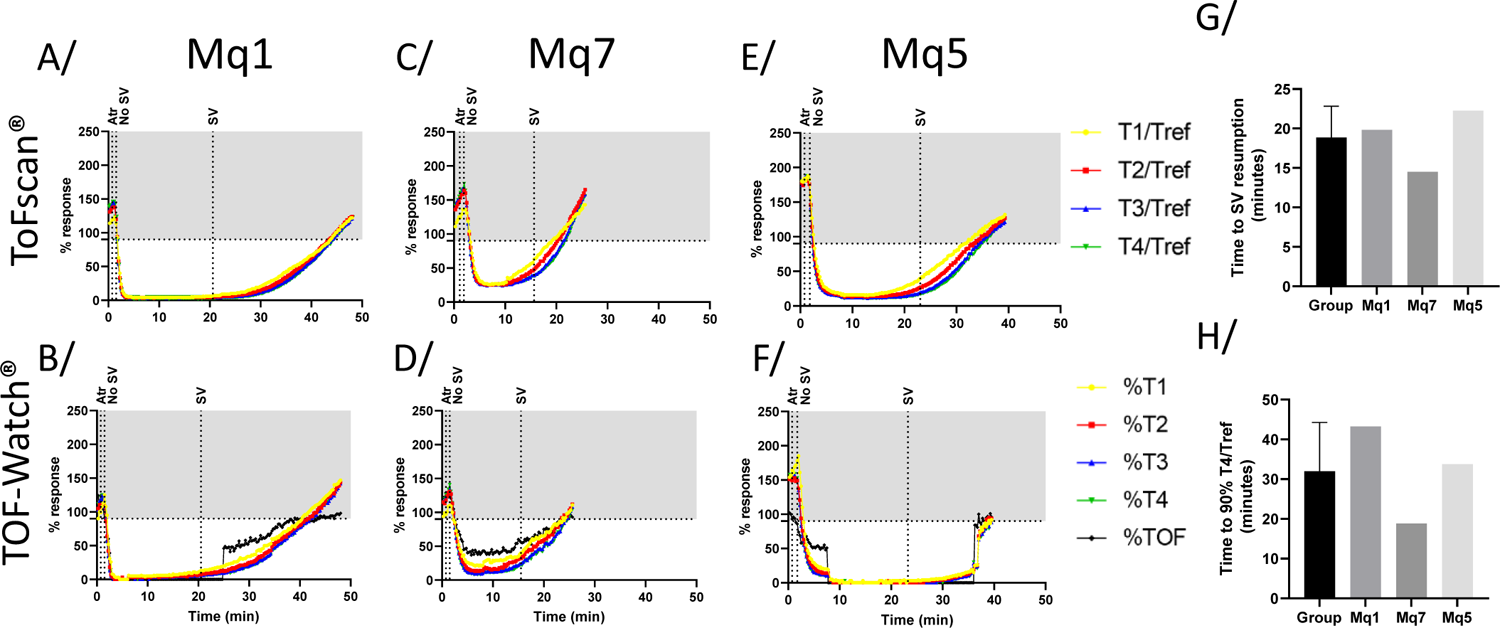
Train-of-four monitoring of three examples of atracurium-enhanced anaesthesia without rescue. The top line (A, C, and E) are follow-ups done with the ToFscan software (IDMED); the bottom line (B, D, F) were done using the TOF-Watch^®^ software (Alsevia). Regarding the Alsevia software, when no %TOF was given, the values were set to 0. Both were coupled with their respective dedicated hardware. Each column represents a single animal (n=3). Atr: atracurium injection, 400µg/kg dose (i.v). No SV: loss of spontaneous ventilation. SV: Spontaneous ventilation resumption.

**Figure 5:**
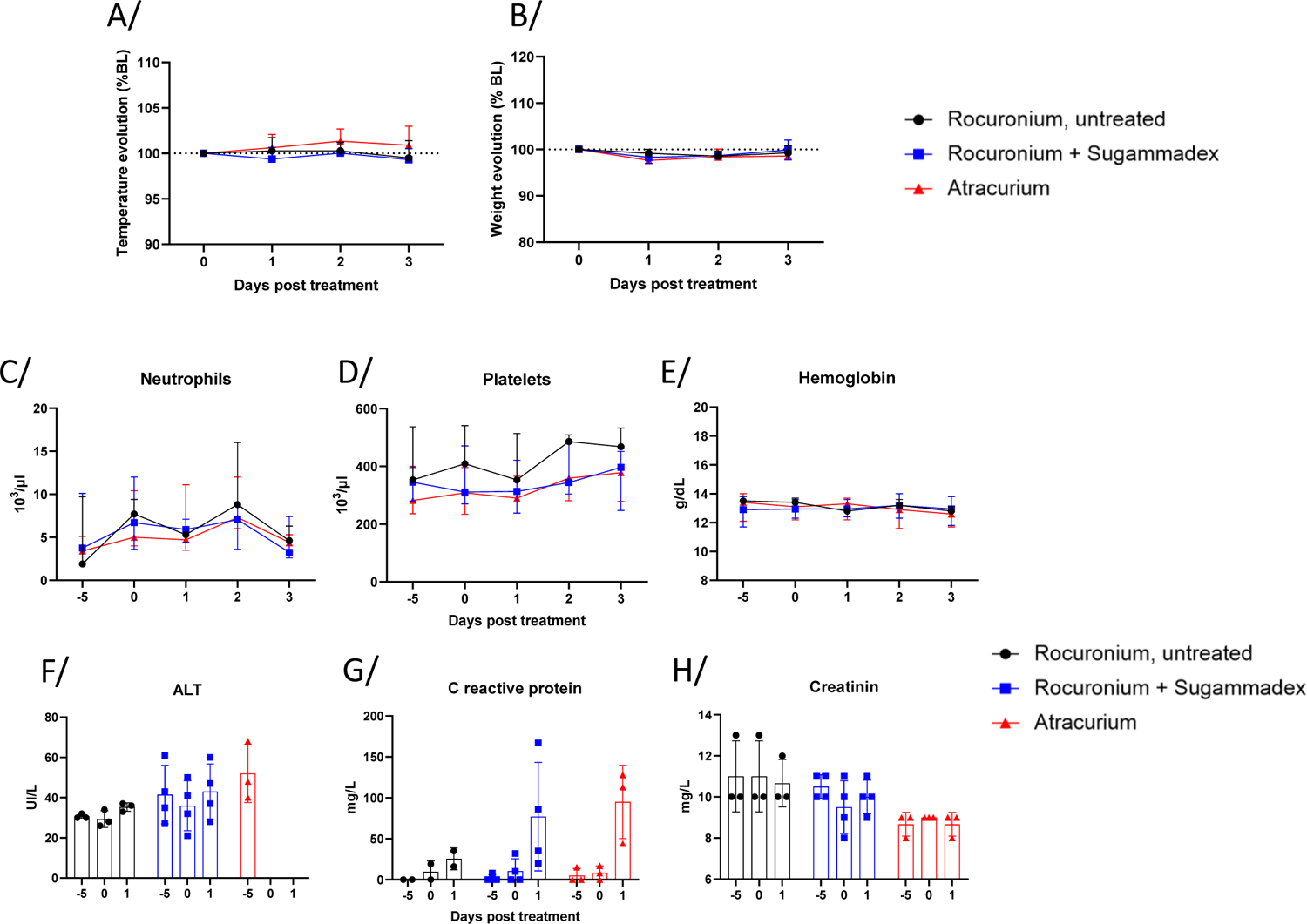
Clinical follow-up of curare enhanced anaesthesia experiencing animals. A-B/ Median and standard deviation of temperature and weight modification reported to the day of curare enhanced anaesthesia. C-E/ Median and standard deviation of relevant hematological parameters in circulating blood. F-H/ Relevant blood biochemistry parameters. ALT= alanine aminotransferase. A blood drawing on day 0 was done before the curare injection. No statistical analyses were positive. Rocuronium n=3; rocuronium + sugammadex n=4; atracurium n=3. One CRP value from an untreated rocuronium animal was excluded due to endogenic pre-existing high levels.

Regarding both of those biological events, some inter-individual variability could be observed (Figure 4 G and H), similarly to humans (see Supplemental). Of note, the delay until full recovery (i.e., TOF ratio > 0.9) appears to be longer with atracurium than with rocuronium but not statistically significant (22.75 minutes versus 33.78 minutes, respectively). Sugammadex did not affect atracurium-induced deep neuromuscular blockade (Supp. Figure 1). Biological measurement was not affected by any procedures (see Supplemental).

## DISCUSSION

NMBA is regularly used in NHP for biomedical research, but only a few data are available on the posology^16,19^, and no on monitoring in macaques. In this study, we were able to assess in cynomolgus macaques the optimal atracurium and rocuronium induction doses to allow a profound neuromuscular blockade that would mimic the pharmacokinetic observed in humans to 0.4 mg.kg^-1^ and 0.2 mg.kg^-1^ respectively. As expected from results obtained in Rhesus monkeys^16,19,20^, sugammadex was able to reverse rocuronium-induced neuromuscular blockade within minutes at a 1 mg.kg^-1^ dose in cynomolgus macaques, with the same kinetic as in humans. No toxicity was observed throughout the procedures.

### NMBA pharmacokinetics in humans

The two non-depolarizing NMBAs, atracurium, and rocuronium, display middle long-lasting effects^8,15^. In humans, atracurium induction dose was established at 0.5 mg.kg^-1^. The loss of the T4 response is usually achieved within 90 to 180 seconds, allowing tracheal intubation, with a complete neuromuscular blockade within 3 to 5 minutes. For rocuronium, the initial dose is 0.6 mg.kg^-1^, allowing tracheal intubation in 90 seconds. Both of them have a clinical effect time (*i.e.,* delay to obtain full recovery) of around 35 minutes (T4/T1 > 90%). In humans, the sugammadex dose to reverse rocuronium-induced blockade depends on the depth of the blockade: 2 mg.kg^-1^ when two twitches are recovered, 4 mg.kg^-1^ when no TOF response is observed or 16 mg.kg^-1^ to stop a complete neuromuscular block immediately after establishment. The objective was to establish the lowest dose of rocuronium and atracurium that could induce a profound neuromuscular blockade within the same time range as in humans (abolishment of 4 twitches within 60-90 seconds).

### Stimulus intensity differences within TOF monitors

Validating the monitoring in macaques was challenging. To be accurate, the decrease in response after NMBA administration has to be caused by the drug itself and not by a decrease in activated fibers or by an increase in skin impedance (underlying the importance of skin preparation), explaining the device calibration recommendation. In adults, calibration is reproducible for both the ToFscan® and the TOF-Watch® at a 50mA intensity. In the experiments we presented on cynomolgus macaques, calibration was possible with the ToFscan^®^ device at a stimulus of 20 mA but not with the TOF-Watch^®^ at any intensity. Nevertheless, we decided to use both TOF monitors simultaneously as we considered they could provide complementary information. The same approach of double-monitoring was evaluated in a paediatric population after rocuronium infusion to compare ToFscan^®^ and TOF-Watch^®^ (the only one that was calibrated), concluding on a good concordance between the two TOF devices, despite the absence of calibration for one of them.^21^ Similar results were reported in an adult population during gynaecologic procedures.^22^

At different stimulation intensity values for ToFscan^®^ (20 mA) and TOF-watch^®^ (10 mA) devices, there were no statistical differences between the two devices to obtain a full recovery (T4/T1>90%). However, they tended to be more delayed with the TOF-watch^®^. In the context of practical use of NMBA and TOF monitoring in this experimental setting, both measurements provided reliable and comparable information: 20 mA with T4/Tref as the evaluation criteria or 10mA with T4/T1 ratio. However, because the muscle-tissue ratio is elevated in monkeys’ upper limb, we believe a 20 mA stimulus might be too important for some animals, possibly leading to a direct stimulation of the macaques’ muscle (when the stimulus applied is greater than the stimulus required for nerve depolarization^15^). Using the ToFscan^®^, the TOF responses never completely vanished during profound neuromuscular blockade, without any fade, supporting the use of the T4/Tref ratio to assess the intensity of the blockade at this intensity. When comparing the two monitoring devices, the absence of T4/T1 response on the TOF-Watch^®^ (10 mA) corresponded to a T4/Tref of 8-10% on the ToFscan^®^ (20 mA), which was the target value range we decided to use herein. Other settings (such as electrode type and placement) were identical and could not explain this difference. The setup worked satisfactorily with great reproducibility, and, considering the anatomical features and relative size of macaques compared to other NHPs, one could propose that NMBA infusions and TOF monitoring could be applied with the same setting to different types of macaques and NHP species, such as the rhesus monkey as reported before.^13,19^

### Anaesthesia consequences on the animals

NMBA-enhanced anaesthesia was of little impact on the animals. CBC and biochemistry were minimally modified, without statistical significance or clinical relevance. Even though a few animals were involved in this study, we found a tendency for higher C reactive protein induction (without haptoglobin) after atracurium or rocuronium+sugammadex than after rocuronium. In experimental models in which acute inflammation monitoring would be required, rocuronium would be the preferred choice. Finally, no adverse events related to sugammadex^23–25^ were observed in our animal cohort.

## CONCLUSION

We were able to assess in cynomolgus macaques the optimal atracurium and rocuronium induction doses to allow a profound neuromuscular blockade that would mimic the pharmacokinetic observed in humans to 0.4 mg.kg^-1^ and 0.2 mg.kg^-1^, respectively. Sugammadex was able to reverse rocuronium-but not atracurium-induced neuromuscular blockade in just over a minute (75 seconds ± 22.8) at a 1 mg.kg^-1^ dose. No adverse event was observed throughout the procedures. As a result, we consider the curare-enhanced anaesthesia methods in NHPs proposed here to be relevant to clinical practice, with minimal impact for different experimental settings, including surgery and mechanically-assisted ventilation. The results and the inter-animal variability reinforce the mandatory use of NMBA monitoring in animals, as currently recommended in human clinical practice.

## ACKNOWLEDGMENTS

We thank Pr Benoit Plaud and Pr Isabelle Constant for their helpful advice and guidance on NMBA monitoring in adults and children.

We are also thankful to Jean-Marie Robert, Maxime Potier, Sébastien Langlois, Thierry Prot, and all ANIMALLIANCE members for carefully handling the animals, and to Sylvie Legendre for biologic measurements.

The Infectious Disease Models and Innovative Therapies research infrastructure (IDMIT) is supported by the “Programme Investissements d’Avenir” (PIA), managed by the ANR under references ANR-11-INBS-0008 and ANR-10-EQPX-02-01.

## AUTHORS’ CONTRIBUTION

AGC: protocol design, animal experiments, data interpretation, writing up of the first draft of the manuscript

FR: protocol design, animals’ selection, animal experiments, revision of the manuscript

JL: protocol design, animals’ selection, animal experiments, revision of the manuscript

EB: animal experiments, revision of the manuscript

HL: protocol design, animals’ selection, animal experiments, data interpretation, writing up of the first draft of the manuscript

RL: protocol design, revision of the manuscript

PB: funding of the study, protocol design, modification of the manuscript

## ABBREVIATIONS

BSL: Bio Safety Level

CEA: Commissariat à l’énergie atomique et aux Energies Alternatives

CRP: C-reactive protein

IV: Intravenous

NHP: Non-Human Primate

NMBA: NeuroMuscular Blocking Agent

rNMB: Residual Neuromuscular Blockade

SV: Spontaneous Ventilation

TOF: Train-Of-Four

