## Supplemental for "Optimization of neuromuscular blockade protocols in cynomolgus macaques: monitoring, doses and antagonism"

### Biological evaluation

Several clinical parameters were assessed before and after curare enhanced anaesthesia. Clinically, no animal exerted signs of discomfort or pain and no behaviour modification could be observed. Furthermore, the animals did not display hyper- or hypothermia, nor significant body weight modification in any of the experimental conditions (Figure 5A-B). Complete blood counts were evaluated in NMBA enhanced anaesthesia with or without reversal before enhanced anaesthesia and 3 days after (Figure 5C-E and Supplementary Figure 1). Neutrophils count (Figure 5C), platelet counts (Figure 5D) and hematocrit levels (Supp. Figure 2) or haemoglobin concentration (Figure 5E) did not vary significantly in either treatment group over the period of observation. No significant variations in WBC, RBC, lymphocytes, monocytes or eosinophils were observed either (Supp. Figure 2). Blood levels of alanine amino-transferase, a marker of liver function, and of creatinine, a marker of kidney function, remained stable whereas the levels of C-reactive protein, a marker of acute inflammation, increased markedly one day after receiving atracurium or rocuronium+sugammadex, (Figure 5F-H). Blood levels of AST, biliary acid, bilirubin, haptoglobin, protein and urea did not vary significantly however (Supplemental Figure 2-3), indicating a good tolerance of atracurium and rocuronium ± sugammadex in macaques.

### NMBA nature and general considerations.

It has been reported that core temperature may affect pharmacodynamics and pharmacokinetics of NMBAs^23^, with lower temperatures reducing twitch intensities and TOF measures. Herein, whereas the warming pad was maintained, the forced-air of the warming blanket was stopped during neuromuscular blockade monitoring, and restarted as soon as the monitoring was ended. The rectal temperature of the animal was therefore monitored continuously and showed no major changes. However, for longer procedure, the warming pad would most certainly be required to keep the animals’ temperature stable and thus adequate monitoring should be verified.

### Animal individual variations.

In two animals, monitoring was particularly challenging to assess with the persistence of four twitches, whatever device was used, even though the muscular contractibility was largely diminished. As reported above, the inter-variability was important within animals, as in humans, supporting the use of monitors to assess the neuromuscular blockade depth and recovery, to ensure proper experiments quality and animal well-being and safety. Loss of the T4 response is usually achieved within 3 minutes after atracurium injection in humans at a 0.5 mg.kg^-1^ dose^24^. When used here at a lower dose in macaques (0.4 mg.kg^-1^), we however observed a consistently faster neuromuscular blockade induction than in humans (median 48 seconds ± 8.4), which is most certainly due, at least in part, to the respectively smaller size of the animal. Such difference should be considered when using atracurium in NHP experimental settings. Median of time needed for spontaneous ventilation (SV) resumption and recovery of 90% TOF ratio did not defer between atracurium and rocuronium (supplemental figure 4).

### Supplemental figure legends

Supplementary figure 1: Train-of-Four monitoring of atracurium enhanced anaesthesia with sugammadex infusion. The top line (A, C) are follow-ups done with with ToFscan® software (IDMED); the bottom line (B, D) were done using the TOF-Watch® software (Alsevia). Regarding the Alsevia software, when no %TOF were given, the values were set to 0. E/ Time (in minutes) needed for T4/Tref resumption from >20% to >90% in the context of a atracurium injection with or without sugammadex infusion. This was calculated using the Idmed software results. Atr: atracurium injection, 400µg/kg dose (i.v). Sug: Sugammadex injection,1 mg/kg dose (i.v). No SV: loss of spontaneous ventilation. SV: Spontaneous ventilation resumption. Both were coupled with their respective dedicated hardware. n=2.

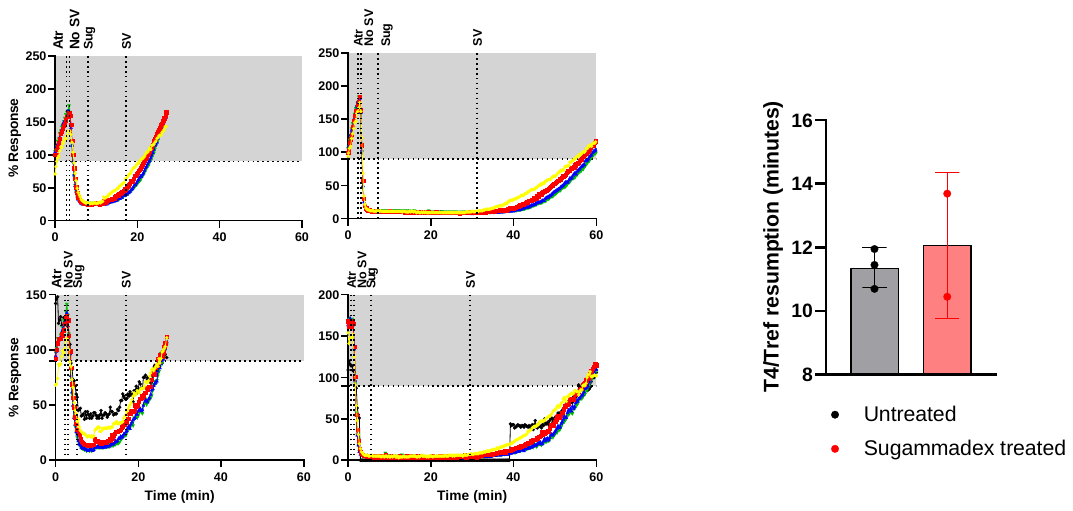

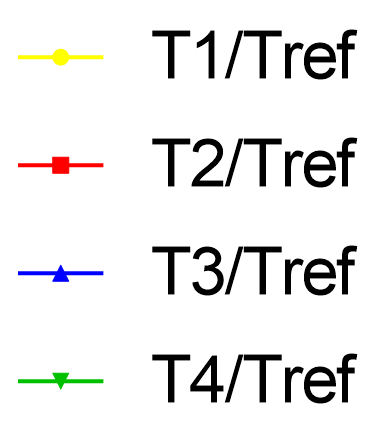

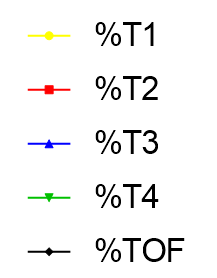

NS

Supplementary figure 2: Additional hematological parameters. No statistical analysis were positive. WBC: white blood cells; RBC: red blood cells. Rocuronium n=3; rocuronium + sugammadex n=4; atracurium n=3.

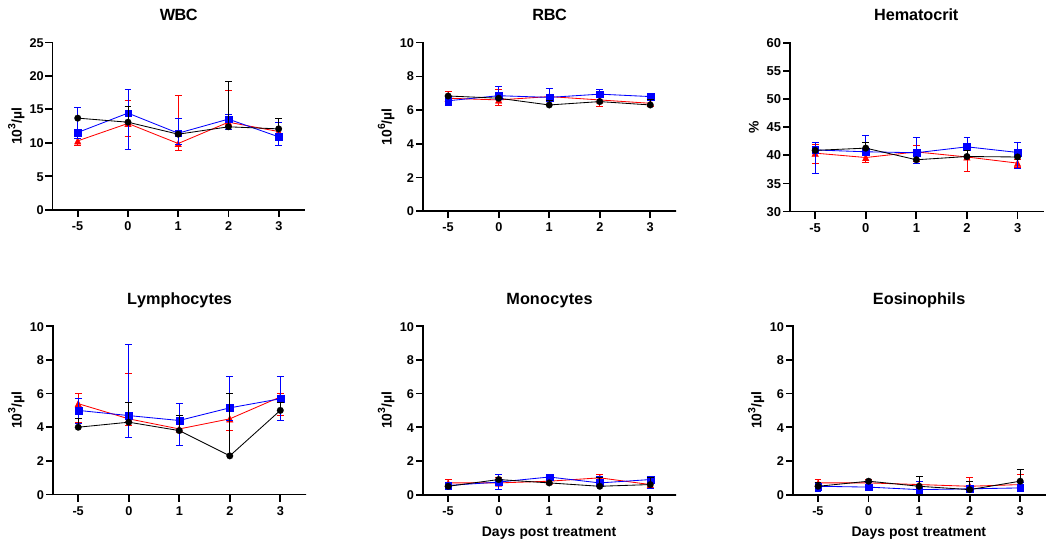

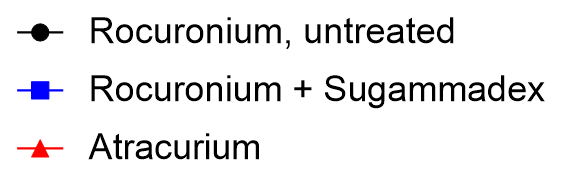

Supplementary figure 3: Additional blood chemistry parameters. No statistical analysis were positive. AST: aspartate amino-transferase. Rocuronium n=3; rocuronium + sugammadex n=4; atracurium n=3.

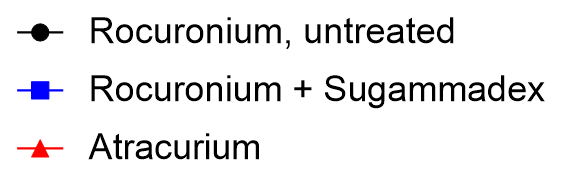

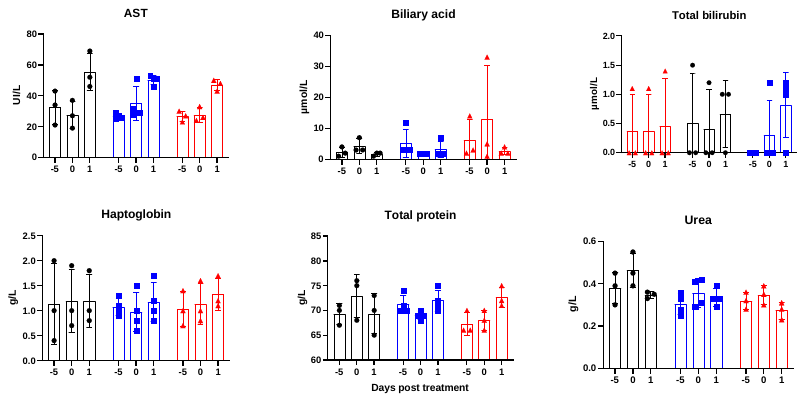

Supplementary figure 4: Direct comparison of rocuronium and atracurium. Median of time needed for spontaneous ventilation (SV) resumption and recovery of 90% TOF ratio. Data were analysed using Mann-whitney non parametric test and were not significant. n=3/group

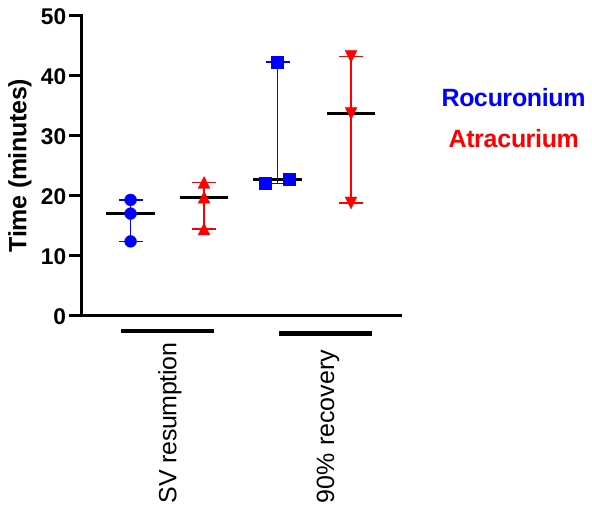
